# High-throughput 5’P sequencing enables the study of degradation-associated ribosome stalls

**DOI:** 10.1101/2020.06.22.165134

**Authors:** Yujie Zhang, Vicent Pelechano

**Affiliations:** SciLifeLab, Department of Microbiology, Tumor and Cell Biology, Karolinska Instituet, Solna, 171 65, Sweden

## Abstract

RNA degradation is critical for gene expression and mRNA quality control. mRNA degradation is connected to the translation process up to the degree that 5’-3’ mRNA degradation follows the las translating ribosome. Here we present an improved high-throughput 5’P degradome RNA sequencing method (HT-5Pseq). HT-5Pseq is easy, scalable and uses affordable duplex-specific nuclease based rRNA depletion. We investigate *in vivo* ribosome stalls focusing on translation termination. By comparing ribosome stalls identified by ribosome profiling, disome-seq and HT-5PSeq we identify that degradation-associated ribosome stalls are often enriched in Arg preceding the stop codon. On the contrary, mRNAs depleted for those stalls use more frequently TAA stop codon preceded by hydrophobic amino acids. Finally, we shown that termination stalls identified by HT-5Pseq, and not by other approaches, are associated to decreased mRNA stability. Our work suggests that ribosome stalls associated to mRNA decay can be easily captured by investigating the 5’P degradome.

## INTRODUCTION

RNA degradation is an integral player of gene expression, as its balance with RNA synthesis is what determines final mRNA abundance. Thus, to understand gene expression, it is important to explore not only mRNA synthesis but also the factors that shape mRNA degradation. Investigating the RNA degradome is important to understand for example differential sensitivity to specialized RNA decay pathways (1) or endonucleolytic cleavage (2). In addition to those specialized degradation mechanism, general processes such as translation are also key to shape the mRNA degradome (3, 4). By investigating 5’phosphorylated (5’P) mRNA degradation intermediates, we have previously demonstrated that in vivo 5’-3’ co-translational degradation provides information regarding ribosome dynamics and codon specific ribosome stalls. Specifically, in budding yeast the 5’-3’ exonuclease Xrn1p follows the last translating ribosome generating an *in vivo* footprint of its position (3). A similar phenomenon has also been described in other organism such as *Arabidopsis thaliana* or rice (*Oryza sativa*) (5, 6). We have previously shown that the *in vivo* ribosome footprint as measured by 5’P degradome sequencing (5PSeq) is a good proxy for general ribosome positions (3, 7). However, as 5PSeq focuses on those ribosome footprints associated with mRNA degradation, in principle it should be enriched in those ribosome stalls leading to mRNA decay. “Ribosome stall” is a broad term that describes increased ribosome density at a particular position, however it does not distinguish between transient pauses and stable stalls (8). General methods such as ribosome profiling investigate the composite of all ribosomes in the cell independent of the functional consequences of those stalls. And even optimized ribosome profiling approaches focusing on disomes, identify ribosome stalls that are not necessarily associated to decay (9). However, It is well known that factors such as codon optimality and ribosome stall often trigger RNA decay (10, 11). All this shows the need for an optimized method able to identify ribosome stalls prone to decay. Nevertheless, despite our ability to investigate the 5’P mRNA degradome, current sequencing approaches are still cumbersome. Methods originally developed to investigate miRNA cleavage such as PARE (parallel analysis of RNA ends) (6, 12, 13) and GMUCT (genome-wide mapping of uncapped and cleaved transcripts) (5) are labour intensive requiring multiple ligation steps, purification and, in some cases, labour-intensive gel size-selection. Even our original 5PSeq protocol requires multiple enzymatic steps and bead-based purifications that limits its application to investigate hundreds of samples (14).

In addition to investigate the presence of 5’P mRNA degradation intermediates, it is important to capture their full diversity. Cytoplasmic mRNA decay in eukaryotes often initiates through deadenylation (15, 16). After poly(A) tail shortening by deadenylase complexes PAN2/3 and CCR4-NOT, mRNA can be degraded exonucleolytically either in 5’-3’ or 3’-5’ orientation. In some cases, 5’-3’ mRNA degradation by XRN1, is independent on deadenylation process (e.g. during NMD) or through the endoribonucleolytic decay (1). This has been confirmed in *Arabidopsis* showing that XRN4 substrates (XRN1 in yeast) can be both polyadenylated and non-polyadenylated. The existing diversity regarding 5’P mRNA degradation intermediates poly(A) length suggests that multiple subpopulations of degradation intermediates coexist. We have previously shown that both oligo-dT and random hexamer priming can be used to identify mRNA degradation intermediates (3, 14). However, identifying both subpopulations in the same experiment and enabling their parallel analysis will offer a more general view of the mRNA degradome.

To facilitate the investigation of the 5’P mRNA degradome in relation with the translation process, we have developed a high throughput 5’P sequencing approach (HT-5Pseq). This approach is flexible, cheap and scalable. We benchmark HT-5Pseq against 5PSeq and investigate its applicability in *S. cerevisiae* and *S. pombe*. We demonstrate it offers high quality degradome information at a fraction of the costs and with significantly decreased hands on time. Additionally, by differentially capturing those partially deadenylated intermediates of degradation, it improves the resolution around the stop codon. We use this approach to compare ribosome stalls at termination level between different growth conditions and show that they are actively regulated. Finally, we investigate the functional consequences of those stalls by comparing HT-5PSeq with ribosome profiling and disome-seq. We focus on those features frequently associated to degradation-associated ribosome stalls and investigate their relationship with mRNA stability.

## MATERIAL AND METHODS

### Growth Conditions and Sample preparations

*Saccharomyces cerevisiae strain* BY4741 (MAT a *his3*Δ*1 leu2*Δ*0 met15*Δ*0 ura3*Δ*0*) was grown to mid exponential phase (OD_600_~0.8) at 30 °C using YPD (1% yeast extract, 2% peptone, 2% glucose). *Schizosaccharomyces pombe* (h-) was grown at 30°C to mid-log phase (OD600~0.8) using YES media (0.5% yeast extract, 3% glucose, supplemented with 225 mg/l of adenine, histidine, leucine, uracil and lysine). For cycloheximide (CHX) treatment, CHX was added to final 0.1 mg/mL to the medium and incubated for 5 min at 30°C for *S. cerevisiae* and 10 min for *S. pombe*. For early exponential phase, strains were grown at 30°C from an initial OD_600_~0.05 to a final OD_600_ of 0.3. To reach early stationary phase, *S. cerevisiae* strains were grown during 60h in YPD. All yeast samples were collected by centrifugation and pellets were frozen in liquid nitrogen. Total RNA was isolated by the standard phenol:chloroform method, and DNA was removed by DNase I treatment. RNA integrity was checked by agarose gel.

### HT-5Pseq library preparation and sequencing

For HT-5Pseq libraries construction, we used 6 μg of DNA-free total RNA. Samples were directly subjected to RNA ligation. The treated RNA samples were incubated with 100 μM RNA rP5_RND oligo (final 10 μM, Table S3) 2h at 25°C with 10 Units of T4 RNA ligase 1 (NEB). Please note that we used an RNA oligo, and not the DNA/ RNA oligonucleotide previously used (14). Ligated RNA was purified with RNA Clean XP (Beckman Coulter), according to the manufacturer’s instructions. RNA was reverse transcribed with Superscript II (Life Technologies) and primed with Illumina PE2 compatible oligos containing random hexamers (20 μM, Table S3) and oligo-dT (0.05 μM, Table S3). Reverse transcription reaction was incubated for 10 min at 25°C, 50 min at 42°C and heat inactivated for 15 min at 70°C. To deplete RNA in RNA/cDNA hybrid after reverse transcription, we used sodium hydroxide (40 mM) for incubation 20 min at 65°C and then neutralized with Tris-HCl, pH =7.0 (40 mM). For DSN (Duplex-specific nuclease) based rRNA depletion, we used a mixture of probes (Table S1) targeting the 18S rDNA, 25S rDNA and 5.8S rDNA. The probes were designed to occupy the whole ribosomal RNA regions with consecutive 25-30 nt long unmodified DNA oligos. The hybridization of probes (2 μM each) with cDNAs were incubated at 68 °C for 2 minutes before adding pre-warmed DSN buffer mix with 1 Units of DSN enzyme (Evrogen). The reaction then performed at 68 °C for 20 minutes. To inactive DSN enzyme, we added 2X DSN stop solution and incubate 10 min at 68 °C. The final PCR amplification was performed using 2X Phusion High-Fidelity PCR Master Mix with HF Buffer (NEB) and final 0.1 μM of PE1.0 and corresponding multiplex PE2.0_MTX (Table S3). The program followed this: 30s 98°C; 15 cycles (20s 98°C; 30s 65°C; 30s 72°C); 7min 72°C. Libraries were size selected using 0.7x-0.9x (v/v) AMpure XP beads (Beckman Coulter) to final length of 200-500 bp and sequenced by NextSeq 500 using 60 sequencing cycles for Read 1 and 15 cycles for Read 2.

### Read preprocessing and analysis

3’-sequencing adaptor trimming was applied to 5 ‘ends of reads using cutadapt V1.16 (http://gensoft.pasteur.fr/docs/cutadapt/1.6/index.html). The 8-nt random barcodes on the 5’ ends of reads were extracted and added to the header of fastq file as the UMI using UMI-tools. Reads were mapped to the reference genome (SGD R64-1-1 for *S. cerevisiae* genome, ASM294v2.20 for *S. pombe* genome) separately by star/2.7.0 (31) with the parameter --alignEndsType Extend5pOfRead1 to exclude soft-clipped bases on the 5’ end. To calculate the fraction of rRNA, tRNA, snRNA snoRNA and mRNA in library compositions, the stepwise alignment was performed by corresponding index generated by star/2.7.0. Duplicated 5’ ends of read introduced by PCR during library preparation were removed based on random barcodes sequences using UMI-tools (32). To compare the differences of 5’P read coverage in DSN based rRNA depletion at gene level, reads per gene were counted using Subread package (featureCounts) (33). mRNA, tRNA, rRNA and snRNA and snoRNA transcripts were counted separately and combined for further analysis. Differential gene expression analysis were performed using the DESeq2 packages from R and Bioconductor (http://www.bioconductor.org/) (34). The threshold for differentially expressed genes were defined as p value < 0.005 and log2(fold-change) > 1. Analysis of 5’ ends positions was performed using *Fivepseq* package (20) (http://pelechanolab.com/software/fivepseq), including relative to start, stop codon and codon specific pausing. Specifically, the unique 5’mRNA reads in biological samples were summed up and normalized to reads per million (rpm). Then the relative position of 5’mRNA reads to all codons of all ORF were summed at each position. Metagene plots were showed as the sum value versus the relative distance from respective codon. All clustering analyses was performed by k-means using Complexheatmap packages from R and Bioconductor (35). Gene Ontology enrichment analysis was performed with ClusterProfiler using Fisher’s exact test (36). Significance for enrichment for particular nucleotides and amino acids at each position were performed using kpLogo (37). Ribosome profiling and Disome-seq data were obtained from Guydosh *et al*. (23) and Meydan *et al*. (25), respectively. Termination pause scores were calculated in each dataset by dividing the peaks from termination versus the upstream 27-nt regions (before the following ribosome queuing). Collision scores were calculated from Meydan *et al*. (25) by dividing termination pause scores in disome versus monosome profiling. Dataset for *S. cerevisiae* tRNA adaptation index, mRNA codon stability index, translation efficiency were obtained from Carneiro *et al*. (26), gene expression level and mRNA half-life were obtained from Xu *et al*. (38), Celik *et al*. (39) and Presnyak *et al*. (10), respectively.

## RESULTS

### Development of HT-5Pseq for investigation of 5’-phosphorylated mRNA degradation intermediates

To enable the high-throughput investigation of cotranslation mRNA degradation and ribosome dynamics, we set to improve our previous 5PSeq approach (14). We identified multiple bottlenecks for its systematic application: the high number of enzymatic reactions, the required DNA purification steps and the difficulty associated to rRNA depletion (that compose the majority of the 5’P RNAs in the cell). By improving those steps, we decreased both the required hands-on time as well as the cost per library. In HT-5Pseq, we first ligate an RNA oligo containing unique molecular identifiers (UMI) to the 5’P ends of total RNA (Fig 1A and Fig S1A). Total ligated RNA is subjected to reverse transcription using a mix of Illumina compatible oligos priming with a random hexamer and oligo-dT. This obviates the necessity of *in vitro* RNA or cDNA fragmentation and improves sequencing coverage around the stop codon region (see below). After the generation of cDNAs containing Illumina compatible adaptors at both sides, we degraded RNA with NaOH. To selectively deplete abundant cDNA complementary to rRNA, we designed a pool of affordable non-modified DNA oligos (Table S1). We annealed the rRNA depletion oligos with the single-stranded cDNA library at 68°C and treated the mix with duplex-specific nuclease (DSN) (23), a strategy that has been previously shown to be useful depleting problematic sequences from RNA-Seq at cDNA stage (18). After this, the rRNA depleted cDNA library is purified and PCR amplified to generate the final sequencing library. By eliminating the need for costly biotin-based rRNA depletion and the time-consuming mRNA fragmentation, we reduced the number of bead-based purifications (from 11 to 4) and eliminated the enzymatic steps required for endrepair, dA-tailing and DNA ligation. All these modifications reduced the total time from 35 h to 9 h and the library cost preparations 70% (Fig S1A and Table S2).

**Figure 1.**
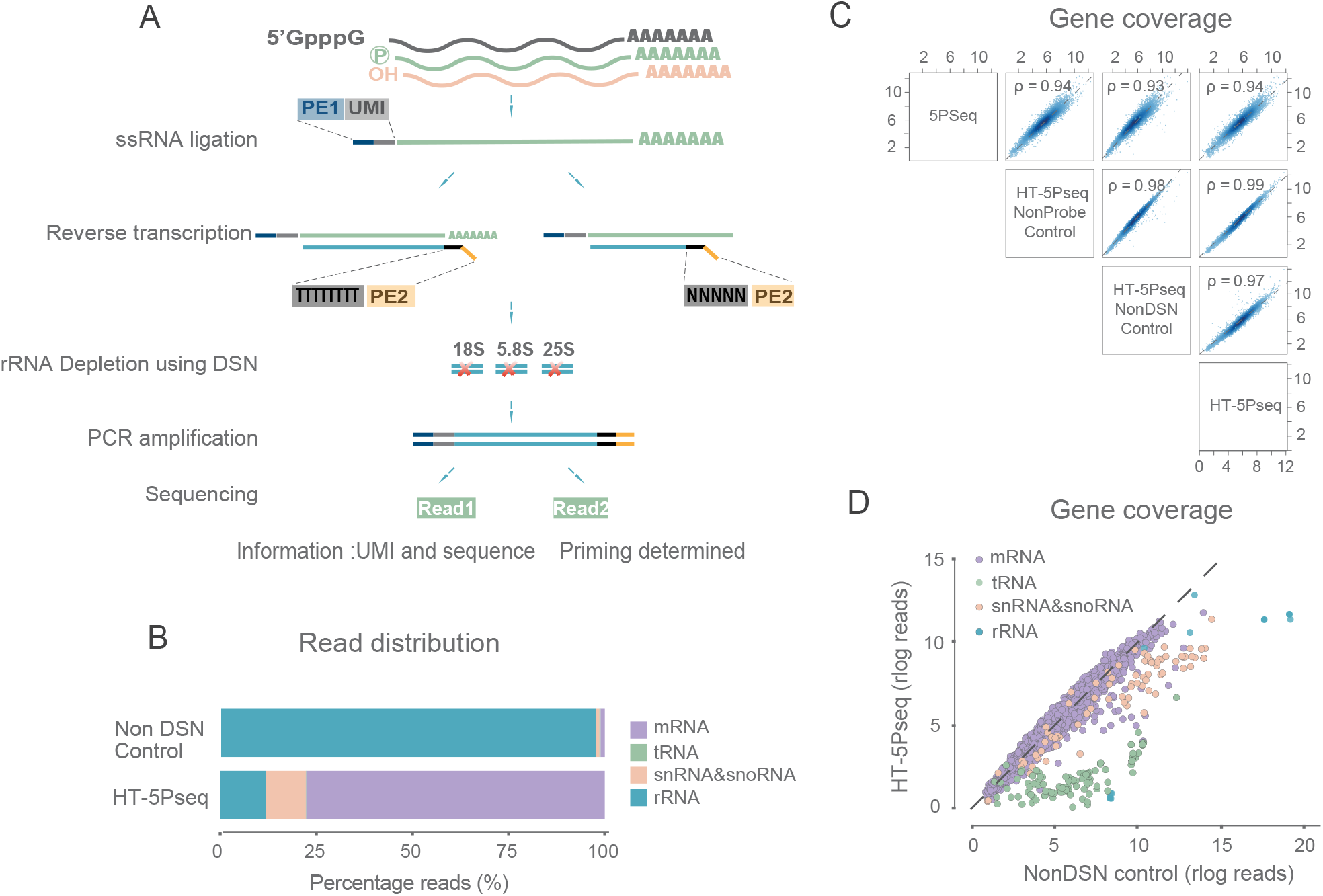
Development of HT-5Pseq. (A) HT-5Pseq method outline. A specific RNA oligo (PE1) is ligated to 5’P RNA molecules. RNA is reverse transcribed using as primer a mix of sequencing oligos (PE2) containing oligo-dT and random hexamers. The cDNA originating from rRNA is depleted using DNA oligos and doublestrand specific nuclease (DSN). cDNA is PCR amplified and sequenced. (B) Improvement of mRNA mapability after rRNA depletion, comparing HT-5Pseq and same protocol omitting DSN treatment (%). (C) Spearman correlation between 5PSeq (3) and HT-5Pseq. NonDSN refers to control libraries omitting DSN rRNA depletion. NonProbe refers to libraries treated with DSN but omitting the depletion oligos. 5’P read gene coverage shown in rlog. (D) Differential gene-specific 5’P read coverage.

DSN-based rRNA depletion increases the percentage of reads aligned to mRNAs from 1.1% to 77.6% (Fig 1B) and also the associated sequencing library complexity (Fig S1B). Accordingly, DSN treatment also decreases the rRNA reads percentage from 98.0% to 12.0% (Fig 1B). Although the main application of HT-5Pseq is the identification of regions with an increased 3-nt periodicity of 5’P fragments (e.g. as those caused by ribosome stalls) and not their overall abundance, we confirmed that DSN treatment is highly reproducible (Fig 1C and Fig S1C-D) and has limited effect on mRNA coverage (Fig 1D). DSN treatment decreases sequencing coverage of the depleted rRNA regions (in blue), and other features with significant secondary structure as tRNAs (in green). Only a few mRNAs (0.57%, 31 of the 5398 detected) present significant decreased coverage, likely caused by their ability to form hairpins at 68°C. An advantage of HT-5Pseq is the simple and flexible design of rRNA depletion probes. Contrary to ribosome profiling, in HT-5Pseq samples are not subject to *in vitro* RNA degradation generating small rRNA fragments (19). In 5’P mRNA degradome sequencing, mature rRNA (and tRNAs) with well-defined 5’P boundaries are the main contaminants. To show the flexibility of our rRNA depletion approach, we used probes originally designed for *S. cerevisiae* (Table S1) to investigate the 5’P degradome in the distant yeast *S. pombe*. We confirmed that this treatment increased the percentage of usable mRNA molecules, from 5.0% unique mRNA degradation intermediates to 62.3% after DSN treatment (Fig S1E-F), even when using a suboptimal rRNA depletion panel. All this confirms that HT-5Pseq is a flexible protocol that can be easily adapted to the organism of interest.

### HT-5Pseq provides single nucleotide resolution insights into 5’-3’co-translational decay

Once confirmed its general applicability, we verified HT-5Pseq ability to identify ribosome stalls at single-nucleotide resolution (3, 14). As expected, HT-5Pseq recovers the characteristic 3-nt periodicity associated to 5’-3’ co-translational mRNA degradation (Fig 2A). To further confirm its sensitivity, we investigated codon specific ribosome stalling using our newly developed computational pipeline *Fivepseq* (20). We clearly identified pauses associated to slow codons like arginine (encoded by CGA, CGG) and proline (encoded by CCG) (Fig 2B-C) as we and other have previously shown (7, 21). Reassuringly, we observed similar codon specific pauses in HT-5PSeq to those observed when the same sample was analysed using our previously developed 5PSeq (Fig 2C).

**Figure 2.**
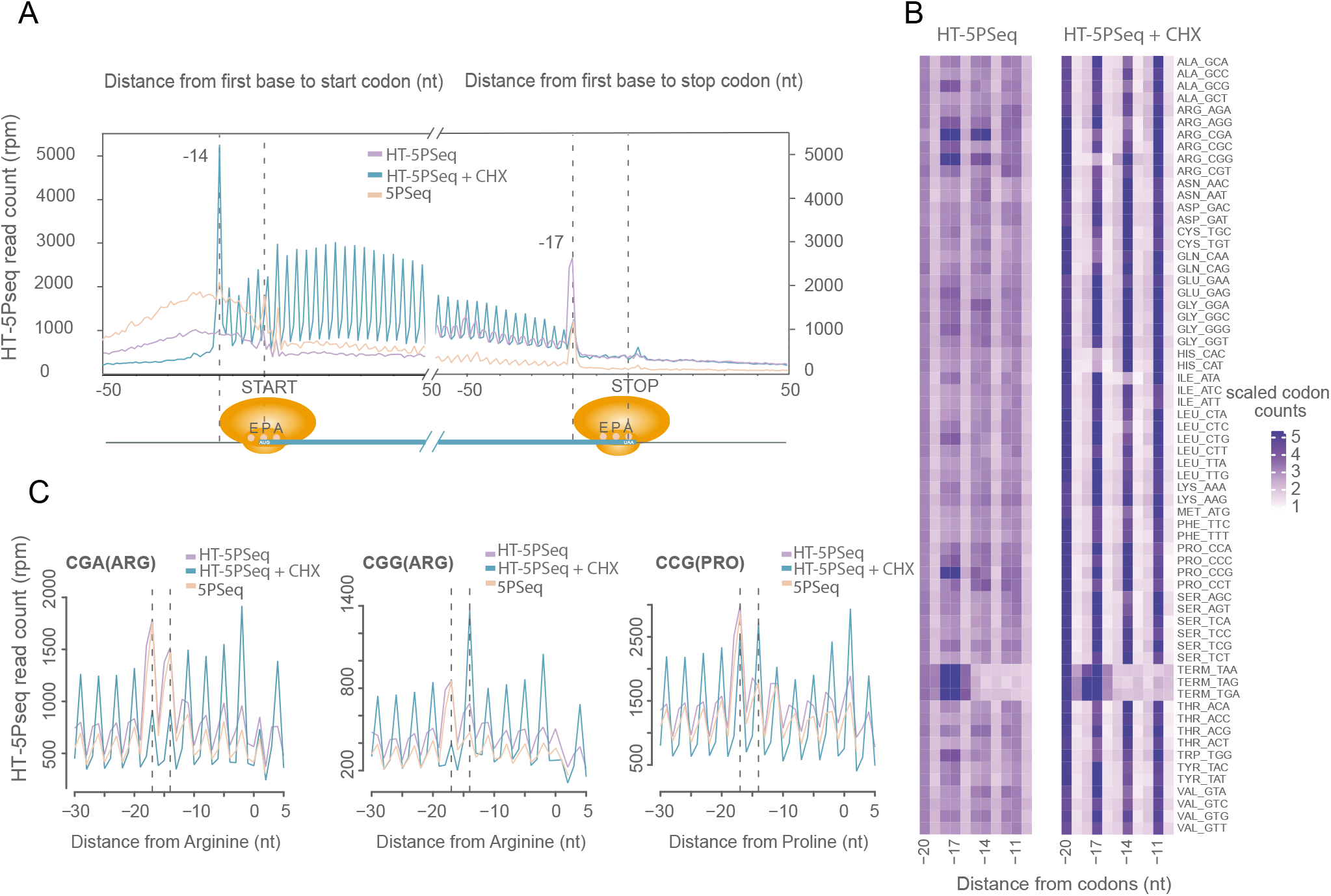
HT-5Pseq reveals ribosome dynamics at codon resolution. (A) Metagene analysis for 5’P read coverage relative to ORF start and stop codon. Cells grown in rich media (YPD) by HT-5Pseq are shown in purple, cycloheximide treated cells (CHX) for 5 mins in blue and 5PSeq are shown in pink. (B) Heat map for codon specific 5’P coverage. Positions −17, −14 and −11 nt represent ribosome protection at the A, P and E site respectively with or without CHX treatment. For each codon, reads were normalized using the total 5’P reads between −30 to 5 nt. (C) 5’P reads coverage for rare arginine (CGA, CGG) and proline codons (CCG). Dotted lines at −17 and −14 corresponding to the expected 5’ end of protected ribosome located at the A site or P site,

To demonstrate the ability of HT-5Pseq to detect direct perturbations of the translation process, we investigated mRNA degradome changes after cycloheximide (CHX) treatment. As expected, CHX clearly increases the observed 3-nt periodicity, especially at the 5’regions of the genes (Fig 2A). In addition to its general effect inhibiting translation elongation, CHX can also mask *in vivo* ribosome pausing at codon level (22). As in ribosome profiling, in HT-5Pseq rare codons present higher 5’P protection indicating ribosome stalling, which was lost after CHX treatment (e.g. correlation between A-site 5’P footprint and tRNA adaptation index (tAI) goes from Pearson correlation r −0.385 to 0.061, Fig S2A). Accordingly, we observe that the protection pattern for proline (CCG) and arginine (CGG) is displaced one codon after CHX treatment (Fig 2C). We also observed an increase protection for codons specifically pausing at A, P or E sites (Fig 2B). To investigate *in vivo* ribosome dynamics in a different species, we treated *S. pombe* with CHX and observed a clear increase of the 3-nt periodicity (Fig S2C), as expected by the general translation inhibition (Fig S2B). As in *S. cerevisiae*, rare codons such as CCG (proline) and CGG (arginine) showed higher enrichment without CHX treatment, but lost enrichment at A site in CHX samples (Fig 2C). We also found displaced CGG (arginine) codon from A site to P site in *S. pombe* (Fig S2D). Our results thus confirm that HT-5Pseq offers codon-specific information of the co-translational mRNA decay process.

### Environmental regulation of ribosome termination pause associated to degradation

An additional advantage of HT-5Pseq is that, by using a combination of random hexamers and oligo-dT primers during reverse transcription, it enables to control the sequencing depth around the stop codon (Fig 3A). In ribosome profiling, coverage around the stop codon can be modest, which can be explained by a combination of biological and technical reasons. For example, footprints associated to translation termination can be lost during the *in vitro* RNA digestion required in ribosome profiling (23, 24), especially when using high salt conditions (21) (Fig S3A), or because the associated footprints possess different length and thus are often excluded from the analysis. In addition, those ribosomes stalls at termination level leading to mRNA decay could be underrepresented in ribosome profiling data (containing the pool of all ribosome footprints), while they should be easy to detect using HT-5Pseq. To further increase the coverage around the stop codon, we focused on those HT-5Pseq reads primed by oligo-dT (Fig 3A). When analysing the sequence coming from read 2 (Fig 1A), we can observe a clear bimodal distribution regarding the number of continuous As sequenced (Fig S3B). As expected, a fraction of reads originates from random hexamer priming (<5s As), while other fraction originates from oligo-dT primed cDNA. Interestingly, in some cases we observed more than 10 As suggesting that oligo-dT priming could also occur downstream on the remaining poly(A) tail. This result also confirms the existence of a fraction of mRNA degradation intermediates containing sizable poly(A) tails. We used the information from read 2 to classify those reads primed by random hexamer (<5s As) or oligo-dT (>9s As). As expected, 5’P degradome reads in the 3’ region of the genes are enriched in oligo-dT primed molecules (Fig 3A). Oligo-dT primed HT-5Pseq reads enabled the observation of a clear protection pattern associated to single (−17nt) and two ribosomes (−47/−50nt, disomes), stalled at the stop codon. This could be observed both at metagene (Fig 3A) and single gene level (Fig 3B). Using gene-specific information, we classified mRNAs according to their relative protection at the stop codon (−17) in respect to the upstream HT-5PSeq coverage as displaying low or high termination pausing (Fig 3B). Next, we investigated the relative expression of genes with high observed termination pause, and found that high expressed genes present an overall decrease of termination pause (Fig S3C). In agreement, mRNAs presenting low termination pauses are enriched for highly expressed GOs related to translation (e.g. cytoplasmic translation (GO:0002181, p-adj < 4.8·10^−9^) (Table S4). On the contrary, those associated with the transcription process had in general increased termination pausing (e.g. DNA-templated transcription (GO:0006351, p-adj < 6.2·10-^3^) (Table S4).

**Figure 3.**
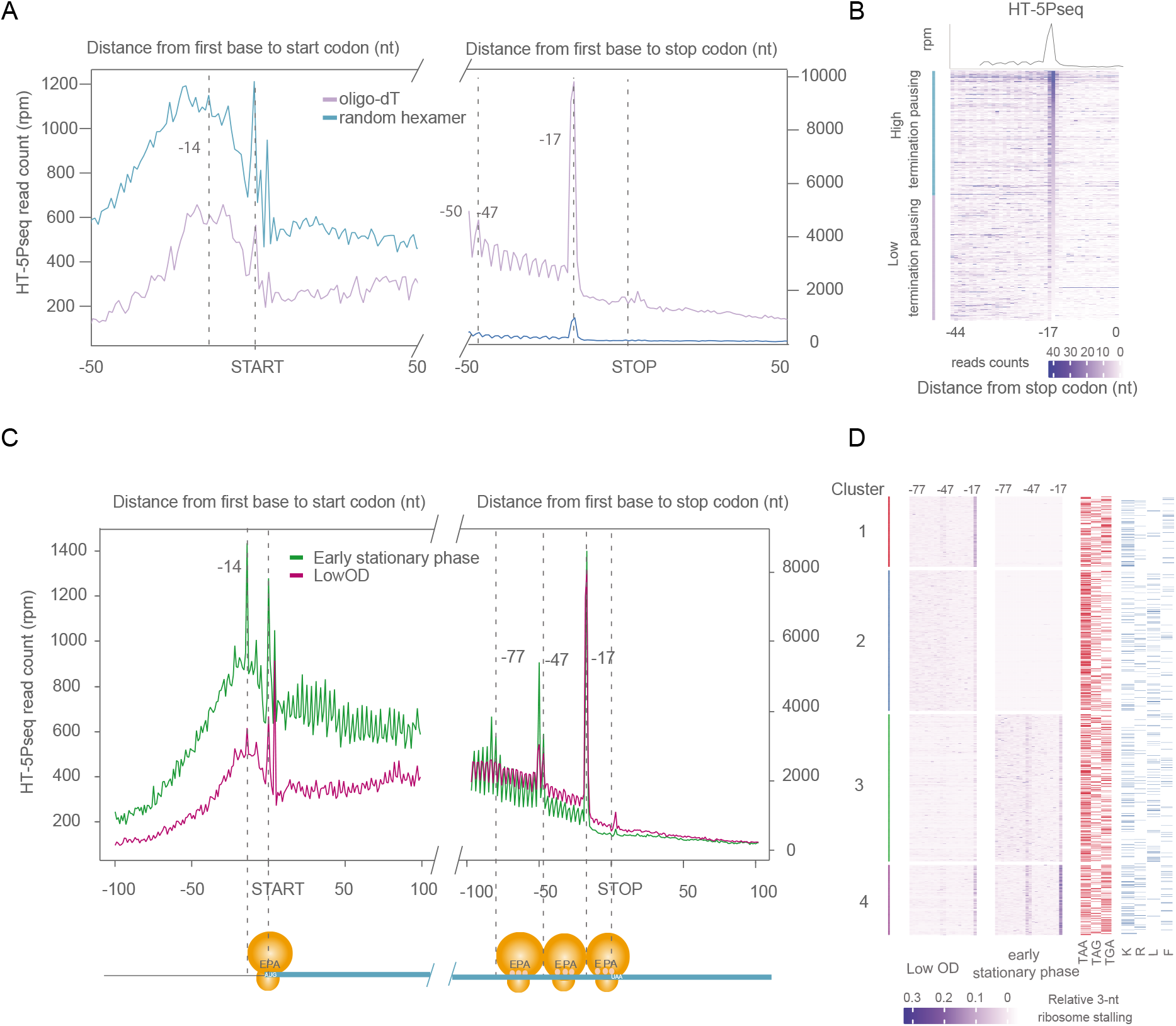
Environmental regulation of termination associated ribosome pauses. (A) Metagene analysis for 5’P read coverage relative to ORF start and stop codon for HT-5Pseq reads primed by oligo-dT(purple) or Random hexamer (blue). (B) Gene-specific 5’P read coverage relative to ORF stop codon. Genes are sorted by the relative ribosome protection at the stop codon (−17) in respect to the upstream region (−44 to −17 nt). Only genes with at least 10 rpm (reads per million) in this region were considered. The top 50% genes were defined as the high termination pausing genes and the bottom half were defined as low termination pausing genes. (C) Metagene analysis for 5’P read coverage relative to ORF start and stop codon. Cells grown at different growth conditions. exponentially growing cells at a lower cell density (OD_600_ ~ 0.3) (in green) and early stationary phase (in purple). (D) Relative 3-nt periodicity for the last 12 codons clustered using K-means. Each point corresponds to the ratio at each codon (ratio between peaks and valleys), not individual 5’P coverage as in B. −17nt indicates a ribosome paused at the stop codon and −50 nt corresponds to disome protection. The stop codon usage is represented in red and amino acids usage before stop codons (lysine, arginine, leucine, phenylalanine) in blue.

Once described degradation associated ribosome stalls at termination level in standard growth condition, we decided to investigate the environmental regulation of this phenomena. We have previously showed that during stationary phase, budding yeast displays massive increase in ribosome stalls at termination level (3). Thus, we decided to compare the degradation associated ribosome stalls in early stationary phase (3 days) and exponentially growing cells at a lower cell density (OD_600_ ~ 0.3). In both cases we observed a clear stop pause at the stop codon (−17). The ribosome pausing of cells at early stationary phase was only limited to −17, but also −47/−50 (corresponding to disomes) and −77/−80 nt (corresponding to trisomes) (Fig 3C). While exponentially growing cells at lower cell density present a very subtle disome peak. Next, we investigated if those pauses were differentially regulated at single gene level. As ribosome pausing was clearly observed at monosome, disome and trisome level (from −80 to −17 nt), we clustered genes according to their relative pausing over the 22 last codons as measured by HT-5Pseq (Fig 3D). We clustered the observed gene specific ribosome pauses in 4 groups. The first 2 clusters had more HT-5Pseq coverage in low OD cells, suggesting a higher gene expression in that condition. Genes in cluster 1 displayed a clear translation termination pausing (−17), while genes in cluster 2 did not. Genes with high translation termination paused in cluster 1 were enriched in RNA metabolic process (GO: 0016070, p-adj < 0.02), while genes with lower translation termination paused in cluster 2 were enriched in cytoplasmic translation (GO:0002181, p-adj < 7.1·10^−30^), similar to what we observed in exponentially growing cells at a higher OD (Table S5). On the contrary, genes in cluster 3 and 4 had more pronounced termination pausing (including disomes) in stationary phase. Genes in cluster 3 are enriched in transcription (GO:0006355, p-adj < 7.5·10^−4^), while genes in cluster 4 were associated with the aerobic respiration (GO:0009060, p-adj < 3.5 ·10^−9^) (Table S5). In summary, our results show that translation termination pausing associated to RNA decay is often environmentally regulated. As ribosome stalling is tightly coupled with mRNA degradation, we decided to investigate up to what degree mRNA degradation, termination pausing, and other mRNA features were related.

### Stop codon context affects termination pausing associated to mRNA decay

To investigate the relationship between ribosome pausing at termination level and co-translational decay, we compared HT-5PSeq with alternative measures of ribosome occupancy such as ribosome profiling or disome-seq (23, 25). We investigated if features associated to termination pauses were shared among those methods that investigate different, although partially overlapping, subsets of ribosome protected mRNAs.

We first defined a simple metric to quantify termination pausing across datasets and investigated paused identified by HT-5PSeq. We quantified the termination pause as the relative protection by a ribosome paused at the stop codon (e.g. 17 nt upstream in the case of HT-5PSeq) and corrected for the for gene specific coverage upstream of that region (see methods for details). After ordering genes according to their level of termination pause, we classified genes with high or low termination pause as we defined previously (Fig 3B). In HT-5PSeq, genes with high termination pause present similar stop codon usage to the genome average (Fig 4A). While genes with lower termination pausing present an increased usage of TAA (65.1%, compared with the high termination pausing of 50.7% and genome average of 47.5 %) and decreased usage of TGA (16.3%, compared with the high termination pausing of 27.5% and genome average of 29.6 %). Observed termination pause was also influenced by the amino acids preceding the stop codon. For example, by the last amino acid preceding the stop codon (Fig 4B). Hydrophobic amino acids preceding the stop codons seem to associate to a decrease termination pausing. Particularly, we found phenylalanine (F) was enriched in genes with low termination pause, as well as other amino acids with hydrophobic chains (e.g. leucine (L) and alanine (A)). While the positively charged arginine (R) was enriched in genes with high termination pause (Fig. S4A).

**Figure 4.**
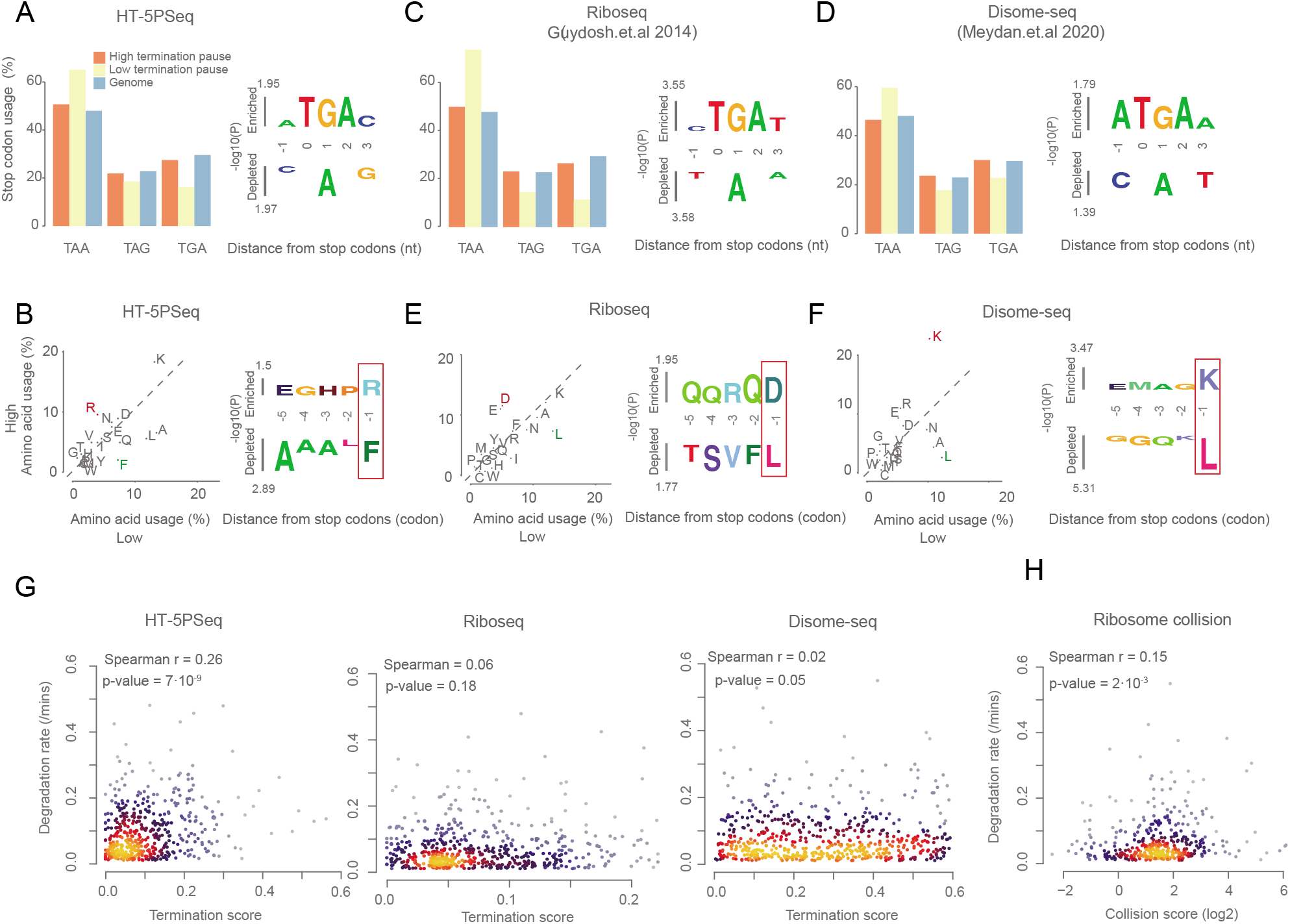
Stop codon context at termination pausing associated to mRNA decay. (A) Frequency of stop codons usage in genes (left) and significance for enrichment for particular nucleotides at each position relative to stop codon (right) using kpLogo (37) by comparing high and low termination pausing groups in HT-5Pseq. (B) Frequency of last amino acid usage in genes with high (y-axis) versus low termination pausing (x-axis) (left) and significance for enrichment for particular amino acid at each position relative to stop codon (right) as in A. (C-D) as A but for ribosome profiling and disome profiling data from Guydosh.et.al (23) and Meydan.et.al (25). (E-F) as B but for ribosome profiling and disome profiling data. (G) Spearman correlation between RNA degradation (calculated by total mRNA half-life from Presnyak et al. (10) with termination pause in HT-5Pseq (left), ribosome profiling (middle) and disome profiling data (right). (H) Spearman correlation between RNA degradation with collision scores. Collision scores were calculated from disome profiling dataset by diving disome peaks with monosome peaks of terminations stalling at stop codons.

To investigate if the same features were associated to general termination pausing, or if they were preferential from degradation-associated ribosome stalls we repeated the same analysis with ribosome profiling and disome profiling data (23, 25). We first compared the measured termination pause between ribosome HT-5PSeq, ribosome profiling and disome-seq. In both ribosome profiling data and disome-seq, the stop codon usage was similar to the one observed in HT-5PSeq (Fig. 4C-D), being mRNAs presenting low termination pause enriched for TAA and depleted for TGA and TAG. This suggest that the degradation-associated ribosome stalls identify by HT-5PSeq share features with stalls identified by methods such as ribosome profiling and disome-seq. Next, we focus on the amino acid context upstream of the stop codon. In ribosome profiling data, which can be expected to represent the global pool of ribosome associated mRNAs, lysine was not enriched in genes with high termination pause. Instead, amino acid with negatively charged sides (i.e. aspartic acid (D)) were enriched in mRNAs with high termination pause (Fig. 4E). However, arginine (R) was enriched before stop codon in mRNAs with low termination pause as in HT-5PSeq (Fig 4B). Then we re-analysed disome profiling datasets that would represent those ribosome stalls long enough to induce ribosome collisions, independent of their ability to induce mRNA decay (9). In disome-seq, we found that lysine (K) was clearly enriched in mRNAs with high termination pause (Fig 4F and S4A). Taking this information together, our results suggest that amino acid-specific preference associated to termination pauses might influence both the degree of termination pausing and their potential resolution and decay. And that degradation-associated ribosome stalls are more similar to those stalls able to induce ribosome collisions.

Finally, we investigated if translation termination pauses as measured by HT-5PSeq were more likely to be associated to mRNA decay than stalls measured by different approaches. Specifically, we investigated the potential association between translation termination pausing and mRNA degradation (10). As expected, we found mild but significant correlation of termination stalling with mRNA degradation rate in HT-5PSeq (spearman cor = 0.26, p-value = 7.0·10^−9^). While the correlation was almost inexistent for general ribosome profiling (spearman cor = 0.06) as well as in disome-seq (spearman cor = 0.02, p-value = 0.05) (Figure 4G). This supported our initial hypothesis that translation termination pause measured by HT-5PSeq are likely enriched in those termination stalls that cannot be resolved and thus lead to increased mRNA decay. To further investigate this phenomenon using an alternative dataset, we calculated “collision scores” of terminations stalling at stop codons by diving disome peaks with monosome peaks from Meydan et.al (9). This measure will reflect the relative enrichment of disome associated stalls in respect to the intrinsic ribosome associated stalls for each gene. As expected, the collision score displayed an increased positive correlation of ribosome collisions with mRNA degradation (spearman cor = 0.15, p-value = 2.0·10^−3^). This suggests that at least part of the translation termination pauses caused by ribosome collision may be associated to decay. To further investigate this process, we analyzed the association of the measured termination pauses (and the collision score) with other features associated to translation and decay (26). Analysing HT-5PSeq data, we found that those mRNAs with lower termination pausing have in general higher codon stability index, codon optimality and translation efficiency (Figure S4B-D). And this effect was even more clear when focusing on the subset of transcripts using TAA as stop codon. In ribosome profiling data, we detected the same tendency as we found in HT-5PSeq. However, this tendency was less clear in disome-seq data, and non-significant when classifying mRNAs according to the collision score (Figure S4B-D). However, it is important to note that the observed decrease could be also explained by the lower sequencing coverage associated to specialized methods such as disome-seq or the interdependence between the ribosome and disome information used to compute the collision scores. Independent of that, our results show that by investigating ribosome termination pauses of different subsets of ribosomes, it is possible to obtain complementary information that can help us to better understand the functional consequences of ribosome stalls.

## DISCUSSION

We have presented here an optimized protocol for the easy measurement of 5’-phosphorylated mRNA degradation intermediates, HT-5Pseq. In this approach, 5’P RNA molecules are first ligated to an Illumina compatible RNA oligonucleotide with UMIs. Ligated RNA is reverse transcribed using a mix of random hexamers and oligo-dT. Contaminant cDNA originating from undesired rRNA molecules is selectively depleted using a pool of affordable DNA oligos and treatment with a duplex-specific nuclease (DSN) (18). This improved protocol decreases hands on time and enables the easy handling of ten to hundred samples, potentially even more if combined with early multiplexing and sample pooling. We have showed that HT-5Pseq is a simple and reproducible approach to detect codon-specific ribosome stalling in both *S. cerevisiae* and *S. pombe*. And that it is specially well suited to investigate those ribosome stalls that frequently lead to RNA decay. We have shown that the use of an oligo-dT primer during reverse transcription increases the coverage at the stop codon. In addition to increase the utility of the approach, this also demonstrates that a fraction of mRNA degradation intermediates still possesses a significant poly(A) tail.

Next, we have investigated the regulation and relevance for mRNA decay of *in vivo* co-translational mRNA degradation footprints associated to ribosome stalling at termination level. It is known that interaction between the ribosome and mRNA sequences nearby stop codon may affect termination pausing. In some cases, this can be attributed to secondary structure downstream of termination pausing (28) or to the interaction of ribosomes with positive lysine amino acid before stop codon, which hinders the binding of the release factors (29). In addition to this, termination stalling can also be enhanced by interfering with the translation termination process, for example, in strains depleted for eIF5A or Rli/ABCE (7, 21, 27). Here we showed the mRNAs presenting lower termination pauses use more frequently TAA as stop codons. To investigate if amino acid usage before stop codon was associated to ribosome stalling, and if this was different between different pools of ribosome-associated mRNAs, we calculated the frequency of amino acid usage in HT-5PSeq, ribosome profiling and disome sequencing. We found that positively charged amino acids (i.e. lysine and arginine) are enriched in HT-5PSeq and disome-seq with high termination pausing. On the opposite, negatively charged amino acids such as aspartic acid is enriched in ribosome profiling with high termination pausing. This suggest that translation termination pausing as measured by the different method measures distinct, but related, biological phenomena. We propose that those differences can be explained by the fact that each method measures a differential subset of ribosomes that differ in the relative ribosome stalling, likehood of ribosome collision and association to co-translational mRNA decay.

Finally, to directly investigate the relationship between co-translational mRNA decay as measured by HT-5PSeq and mRNA stability, we compared ribosome pausing at stop codons with mRNA degradation rates. As HT5-PSeq focus on those mRNAs undergoing degradation, we hypothesized it would be especially useful to identify those ribosome stalls associated to mRNA decay. As expected, we found significantly positive correlation between ribosome stalls measured by HT-5Pseq and mRNA degradation rates. However, that was not the case for ribosome profiling or disome sequencing data. This suggests that those stalls identified by HT-5PSeq are more likely associated to mRNA decay, than those observed by different approaches. Reassuringly, when we calculated a ribosome collision score based on the relative enrichment for disome stalls in respect to intrinsic ribosome (monosome) stalls, we also observed a mild correlation with mRNA decay. This suggests that a fraction of ribosome collisions at stop codons may not be resolved by release factors and undergo decay. However, the functional consequence of those stalls could not be observed when investigation disome or monosome pauses independently. As HT-5Pseq focuses exclusively on the subset of ribosomes undergoing co-translational degradation, we think it offers complementary information to general approaches like ribosome profiling. By using the *in vivo* toeprinting activity of Xrn1, HT-5Pseq can easily capture disomes or trisomes *in vivo*. Although disomes can also be captured using specialized ribosome profiling approaches, such as Disome-seq (28, 29), depending on the stringency of the used *in vitro* RNase I treatment, ribosome footprints can be altered (30). This can lead to the loss of some disome complexes (e.g. those separated by an extra codon) (29). Additionally, as it has been reported non all disome events lead to mRNA decay (25), and thus disome profiling offers different biological information.

In summary, we think that HT-5Pseq will be a valuable tool facilitating the investigation of mRNA life from translation to decay. And that it will be useful to functionally characterize those ribosome stalls leading to mRNA decay. We expect that, by combining multiple genomic and structural approaches, in the future we will be able to understand better this fundamental process in biology.

## ACCESSION NUMBERS

The raw and processed sequencing data are deposited at GEO with accession number GSE152375.

## ACKNOWLEDGEMENT

We thank all members of the Pelechano, Kutter and Friedländer laboratories for useful discussions. We kindly thank Lilit Nersisyan and Jingwen Wang for bioinformatic support. Computational analysis was performed on resources provided by SNIC through Uppsala Multidisciplinary Center for Advanced Computational Science (UPPMAX).

