## Supplementary Material for "High-throughput 5’P sequencing enables the study of degradation-associated ribosome stalls"

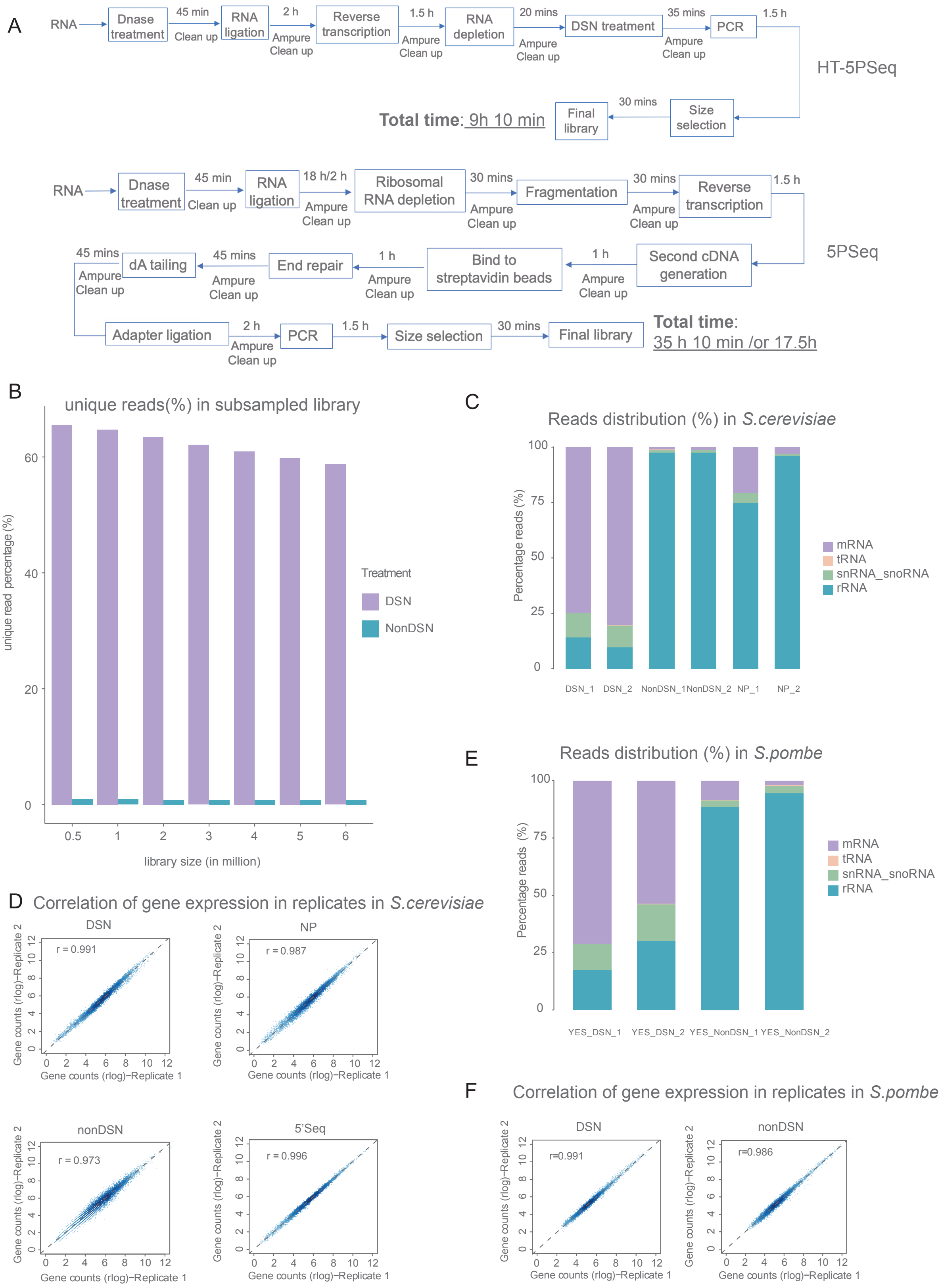

**Figure S1. Performance and reproducibility of HT-5Pseq.** (A) Flowchart of HT-5Pseq and 5PSeq with time estimate for each step. (B) mRNA sequencing library complexity analysis subsampling libraries for HT-5Pseq and control omitting rRNA depletion (NonDSN). Error bars indicate standard errors of the two replicates libraries. (C) Improvement of mRNA mapability in *S.cerevisiae* HT-5Pseq after rRNA depletion. NonDSN refers to control libraries omitting DSN rRNA depletion. NonProbe refers to libraries treated with DSN but omitting the depletion oligos. 2 biological replicates are shown. (D) Spearman correlation between biological replicates for HT-5Pseq, NonProbe, NonDSN and traditional 5PSeq. 5PSeq (3) and HT-5Pseq. NonDSN refers to control libraries omitting DSN rRNA depletion. NonProbe refers to libraries treated with DSN but omitting the depletion oligos. 5´P read gene coverage shown in rlog. (E-F) Differential gene-specific 5´P read coverage. (E and F) as C and D but for *S. pombe.*

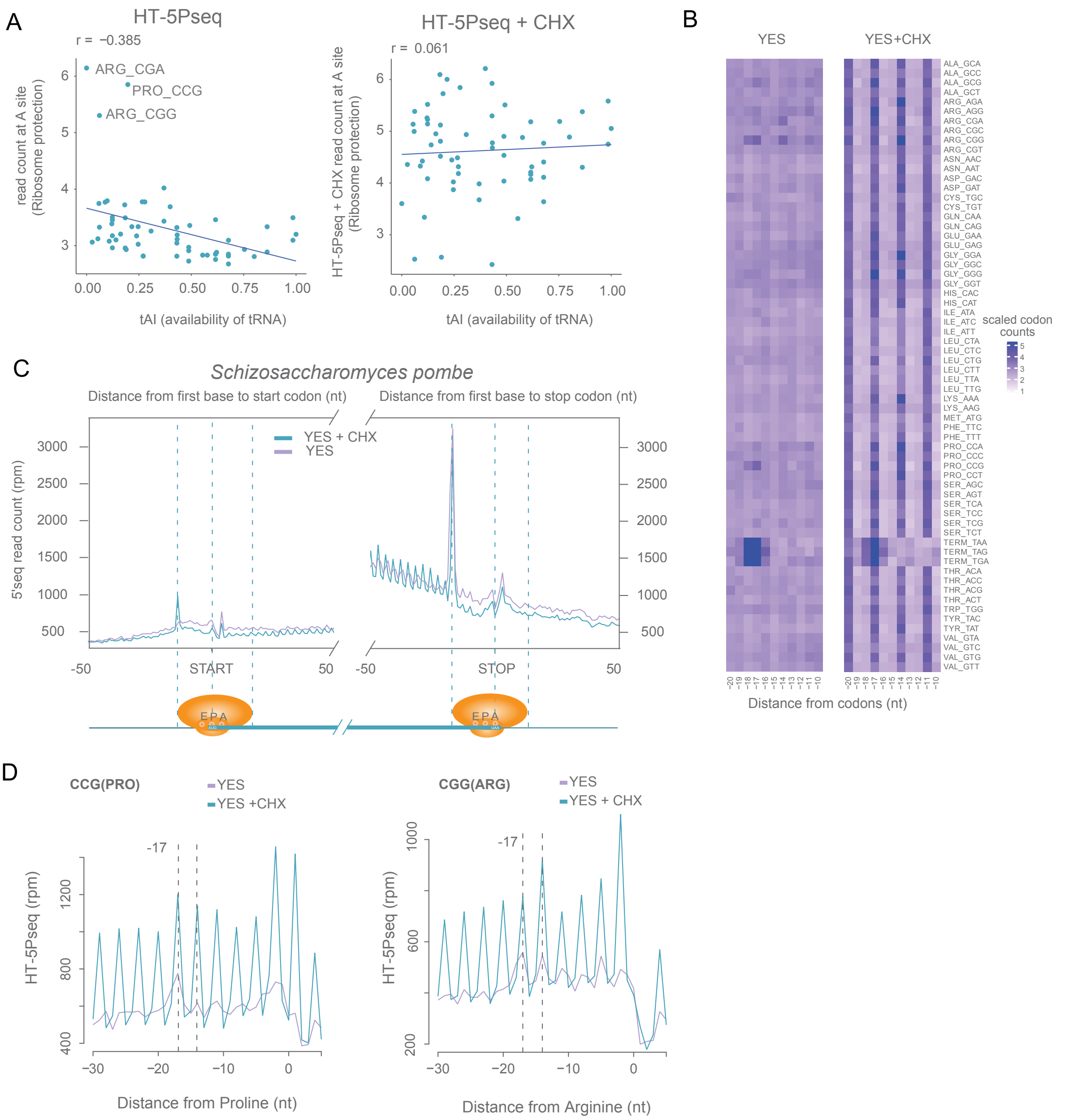


**FigureS2. HT-5Pseq reveals ribosome dynamics in *S. pombe.*** (A) Correlation of 5´P read abundance at A site with tAI (tRNA adaptation index) with or without CHX for *S.cerevisiae.* (B) Heatmap of average codon coverage for cells grown in rich media (YES) and CHX treatment (YES + CHX) in *S.pombe*. 5´P reads relative to each codon were summed up and normalized to the reads corresponding to the surrounding (-30 to 5 nt). -17, -14 and -11 represent A site, P site and E site on ribosome. (C). Metagene analysis showing the abundance of 5´P reads relative to ORF start codons and stop codons for wild type in rich media (YES, in purple) and CHX treatment for 10 mins (YES + CHX, in blue) in *S.pombe*. (D) 5´P reads coverage for proline codons (CCG) and rare arginine (CGG). Dotted lines at -17 and -14 corresponding to the expected 5´ end of protected ribosome located at the A site or P site, respectively in *S. pombe*.

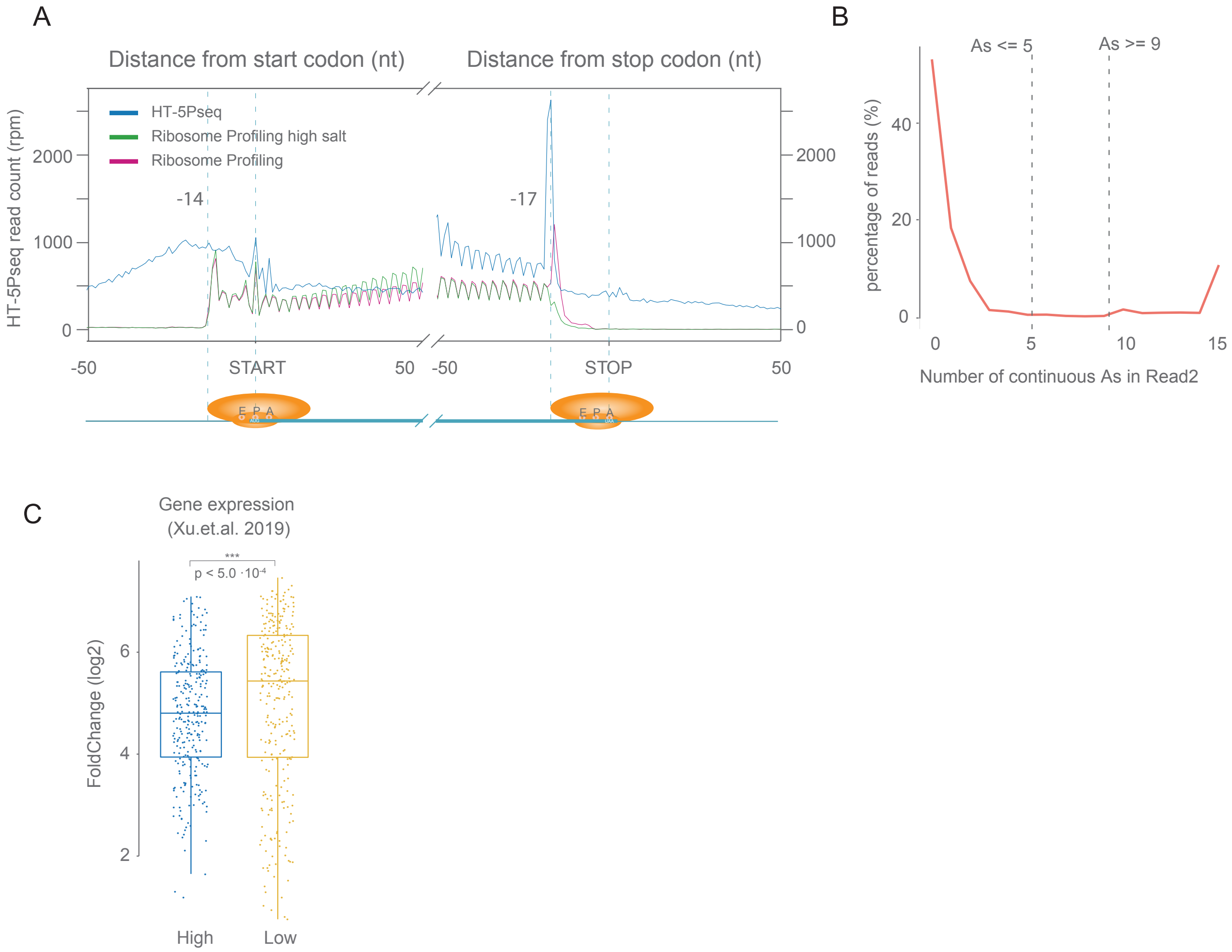


**Figure S3. Influence of stop codon environment on translation termination pauses.** (A) Metagene analysis showing the abundance of 5´P reads relative to ORF start codons and stop codons in HT-5Pseq (in blue), ribosome profiling (in green) and ribosome profiling with high salt wash in lysis buffer (in pink). Ribosome profiling data from Schuller et al (21). (B) The distribution of 5´P mRNA degradation intermediates with different number of continuous As in Read1. (C) Boxplot for gene expression level (array intensity). Gene expression data was obtained from Xu.et.al (38). High termination pause genes are shown in blue and low termination pause in yellow. P-values were calculated by the Wilcoxon Rank Test (two-tailed test).

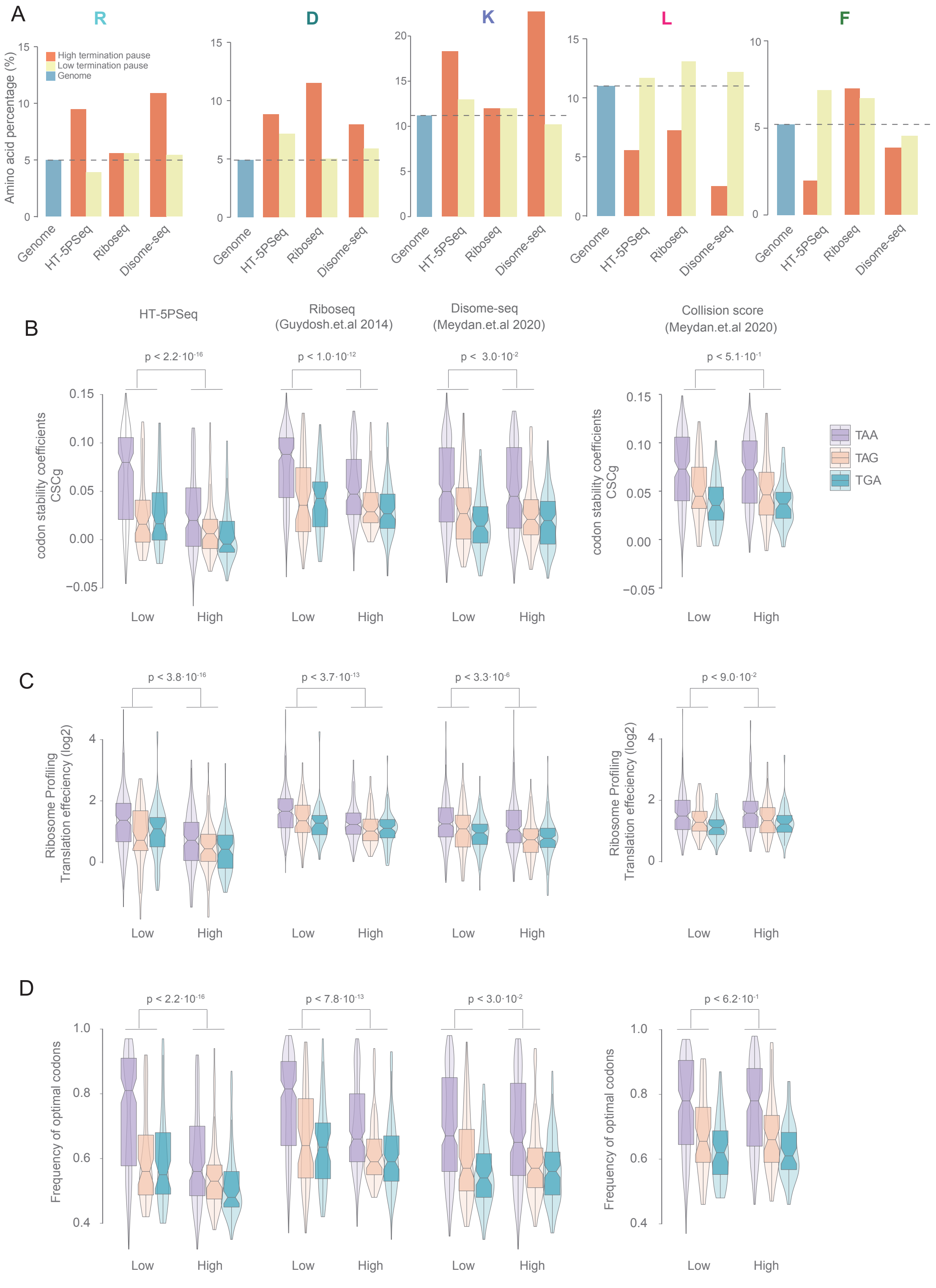


**Figure S4. Characteristics of genes with differential ribosome termination pausing.** (A) Frequency of last amino acid usage (R, D, K, L, F) by comparing genes with high (in orange) and low termination (in yellow) pausing groups in HT-5Pseq, ribosome profiling and disome profiling data. Frequency of last amino acid usage across the genome is shown in blue. (B-D) Comparison of high and low termination groups in: (B) codon stability coefficients (CSCg), (C) frequency of optimal codons and (D) ribosome profiling translation efficiency from Carneiro et al. (26). High and low termination pauses measured as in described in figure 4 for HT-5PSeq, ribosome profiling (23), disome profiling and ribosome collision (25). P-values were calculated by the Wilcoxon Rank Test (two-tailed test).

**Supplementary Table S1.** rRNA depletion oligos for *S. cerevisiae*.

**Supplementary Table S2.** HT-5PSeq library cost estimates.

**Supplementary Table S3.** Oligonucleotides used in HT-5PSeq.

**Supplementary Table S4.** Genes and gene ontologies associated with high and low termination pauses.

**Supplementary Table S5.** Genes and gene ontologies associated with high and low termination pauses during growth condition.
